# Cardiomyopathy-Associated and Basic Residue Mutations in Myopalladin Alter Actin Binding, Bundling, and Structural Stability

**DOI:** 10.1101/2025.09.18.677238

**Authors:** Asha Rankoth Arachchige, Julie Tran, Ziwei Zhao, Hannah Dammann, Alia Michaelis, Vinay K. Kadarla, Michal Zolkiewski, Erika R Geisbrecht, Moriah R. Beck

**Affiliations:** Department of Chemistry and Biochemistry, Wichita State University, Wichita, Kansas, USA; Department of Biochemistry and Molecular Biophysics, Kansas State University, Manhattan, Kansas, USA

**Keywords:** cardiomyopathy, myopalladin, actin binding protein, Ig-domain, sarcomeric protein

## Abstract

**BACKGROUND:** Myopalladin (MYPN) is a striated muscle-specific sarcomeric protein belonging to the immunoglobulin (Ig)-domain-containing family of actin regulators. The Ig3 domain of MYPN is indispensable for interacting with and stabilizing actin filaments, and an increasing number of cardiomyopathic (CM)-associated mutations have been identified within this domain, highlighting its importance for normal cardiac function. Despite their clinical significance, the molecular basis of how these variants disrupt MYPN activity remains poorly understood.

**METHODS:** To identify actin-binding sites, we introduced charge-neutralizing alanine substitutions at basic residues in the Ig3 domain and assessed their impact using actin co-sedimentation assays. CM mutations were similarly evaluated for their effect on actin interaction. Wild-type (WT) and mutant MYPN were expressed in *Drosophila* cardiomyocytes and body wall muscles to assess localization and interaction with actin. The impact of these mutations on protein secondary structure and stability was determined by circular dichroism spectroscopy. In addition, the WT Ig3 domain was further analyzed in co-sedimentation and co-polymerization assays to quantify polymerization of monomeric actin, subsequent bundling and to assess how calcium and salt concentrations modulate MYPN Ig3-actin interactions.

**RESULTS:** MYPN-actin binding is electrostatically driven and mediated by conserved basic charge clusters within its Ig3 domain, a mechanism common among many actin-binding proteins. CM mutations disrupted the MYPN-actin interaction, leading to significant loss of actin bundling without broadly altering secondary structure, except in the destabilized P961L variant. Immunostaining and green fluorescent protein (GFP)-tagged constructs revealed that R955W and P961L mutants accumulated into clusters at the Z-discs. WT Ig3 domain assays further showed that MYPN promotes actin polymerization and crosslinking even under non-polymerizing conditions, and actin-binding activity was not modulated by calcium.

**CONCLUSIONS:** These findings demonstrate that CM-linked MYPN Ig3 mutations compromise actin binding and organization, suggesting that disruption of the MYPN–actin interface is a key pathogenic mechanism in cardiomyopathy.

**What Is Known?:**

- Myopalladin (MYPN) is a striated muscle-specific sarcomeric protein.
- MYPN plays an important role in controlling skeletal muscle growth by regulating actin dynamics and modulating stress-responsive signal transduction and gene expression.
- Mutations in MYPN are linked to various inherited cardiomyopathies (CMs), including dilated, hypertrophic, restrictive, and left ventricular noncompaction forms.

**What new information does this article contribute?:**

- Two conserved clusters of basic residues on the surface of Ig3 domain were identified as key mediators of actin binding and bundling.
- Several cardiomyopathy (CM)-associated MYPN mutations localize to this region, where they impair actin interactions and nearly abolish actin bundling activity.
- Mutants such as R955W and P961L mislocalize abnormally in *Drosophila* cardiomyocytes and body wall muscles.

**Novelty and Significance:** The importance of the *MYPN* gene, which encodes the sarcomeric protein myopalladin (MYPN), was highlighted when 66 mutations in this gene were linked to cardiomyopathies (CMs). MYPN bundles and stabilizes actin filaments and is essential for the organization and maintenance of Z-disc integrity. The C-terminal immunoglobulin (Ig) domains mediate actin interactions, with the Ig3 domain representing the minimal actin-binding unit. Notably, many disease-associated mutations cluster within this actin-binding region and give rise to diverse clinical phenotypes. Despite its clinical relevance, the molecular mechanisms by which different MYPN mutations lead to CM remain poorly understood. Prior mechanistic studies focused mainly on mutations within the cardiac ankyrin repeat protein (CARP) interaction domain of MYPN, which disrupt signaling and nuclear shuttling. In contrast, our study addresses mutations localized to the actin-binding region of MYPN. The effects of these mutations on MYPN-actin interactions had not been previously explored. We show that CM-associated mutations within the Ig3 domain disrupt MYPN-actin interactions, reduce or abolish actin bundling, and induce abnormal subcellular localization patterns in *Drosophila* cardiac and skeletal muscles. This study provides the first direct evidence that disruption of MYPN-actin interactions represents a distinct and clinically relevant mechanism of cardiomyopathy.

## INTRODUCTION

Cardiomyopathies (CM) are a heterogeneous group of myocardial disorders that impair cardiac function^1–4^ and represent a major global public health concern, affecting 6.7 million individuals in the U.S. alone.^1,5^ ^6^ In 2022, CM accounted for 13.9% of all U.S. deaths, making them a leading cause of mortality.^7^ These disorders can be classified as hypertrophic (HCM), dilated (DCM), restrictive (RCM), arrhythmogenic, or left ventricular noncompaction (LVNC) cardiomyopathies and can be further classified as genetic or acquired forms.^4,8^

Mutations in the *MYPN* gene, encoding the sarcomeric protein myopalladin (MYPN), are relatively common among genetic cardiomyopathies.^8^ MYPN is a 145 kDa protein expressed specifically in striated muscle, where it acts as a scaffold linking regulatory proteins in the I-band with structural proteins in the Z-disk assembly while also shuttling to the nucleus to regulate gene expression. A member of the palladin/myopalladin/myotilin family, MYPN shares 68% sequence identity with palladin (PALLD).^4,9–11^ Like PALLD, MYPN contains five immunoglobulin (Ig) domains and a proline-rich region; however, MYPN and PALLD produce opposing effects on actin dynamics. PALLD promotes polymerization, while MYPN inhibits it, though MYPN more strongly stabilizes filamentous (F)-actin against depolymerization.^12^

Among the 66 known MYPN missense variants associated with cardiomyopathies, six (F954L, R955Q, R955W, P961L, C1002W, and R1042C) are localized within the actin-binding Ig3 domain.^13^ Some variants, like P961L, disrupt sarcomere integrity, while others such as R955W preserved structure.^14^ Despite the clinical significance of MYPN mutations in cardiomyopathies, the role of MYPN in cardiac structure and function, as well as the molecular pathophysiology underlying MYPN-associated phenotypic heterogeneity remains poorly understood.^8,13^ Understanding how MYPN maintains Z-disc integrity in muscle tissue may provide insight into therapeutic strategies for MYPN-linked cardiomyopathies.

In this study, we investigate the actin-binding specificity of the MYPN Ig3 domain using charge-neutralizing mutagenesis and actin co-sedimentation assays. Our data reveal that F-actin binding is mediated by conserved basic charge clusters within the Ig3 domain. Given that several cardiomyopathy-associated MYPN mutations localize to this region, we hypothesized that these pathogenic variants disrupt actin interactions and consequently impair sarcomeric integrity, contributing to disease pathogenesis. While the effects of these mutations on MYPN-actin interactions had not been previously examined, our findings show that pathogenic variants impair F-actin binding. Furthermore, the DCM-associated P961L mutation prevented stable Ig fold adoption, while other mutations maintained the overall structure but altered the domain’s stability. To assess the functional consequences *in vivo,* we expressed wild-type (WT) and two DCM mutants, R955W and P961L, in *Drosophila* cardiomyocytes and body wall muscles, revealing distinct subcellular mislocalization patterns relative to Z-disc markers.

Additionally, we found that electrostatic interactions contribute to MYPN Ig3-F-actin binding and that the Ig3 domain functions as an actin polymerizing and bundling protein, even under non-polymerizing conditions. Together, these data establish a mechanistic link between MYPN mutations, defective actin binding, and potential sarcomere disorganization in CM.

## MATERIALS AND METHODS

The corresponding author had full access to all the data in the study and takes responsibility for its integrity and the data analysis. The data that support the findings of this study are available from the corresponding author upon reasonable request. Detailed descriptions of experimental methods are provided in the Supplemental Material.

## RESULTS

### F-actin Binding and Bundling Are Impaired by Neutral Substitutions at Conserved Basic Residues

Previous studies identified two surface-exposed basic patches in the PALLD Ig3 domain (K_13_-LKHYK_18_ and K_51_), along with key lysine residues K36 and K46.^15,16^ Lysine 38, positioned adjacent to K51 in the tertiary structure, also contributes to the second basic patch and has been shown to regulate actin polymerization and bundling.^17^ Sequence alignment of PALLD Ig3 with MYPN Ig3 reveal equivalent basic residues in MYPN (Figure 1A). In the AlphaFold3-predicted structure, residues K949, R950, K952, and R955 cluster into a basic patch on the surface (site 1), while K987 and R988 form a second patch, (site 2)^18^ (Figure 1B). This conservation suggests that basic surfaces may represent a shared, evolutionary conserved actin-binding mechanism. Therefore, we hypothesized that MYPN Ig3 also binds F-actin directly via these basic residues. To test this, we generated alanine substitution mutants at conserved basic positions. Co-sedimentation with F-actin revealed that substitutions at K949, R950, K952, R955, K987, and R988 all reduced F-actin binding affinity. This was evidenced by an increase in the apparent dissociation constant (K_d_) (Figure 2A and Table S1), although the differences were not statistically significant (p < 0.05). While no single mutation completely abolished F-actin binding, all mutants significantly lost the ability to cross-link or bundle F-actin (p < 0.05) (Figure 2B), mirroring results from analogous PALLD Ig3 mutations.^15^ Together, these results support the presence of two conserved actin-binding surfaces in the MYPN Ig3 domain, underscoring their functional importance in cytoskeletal organization.

**Figure 1.**
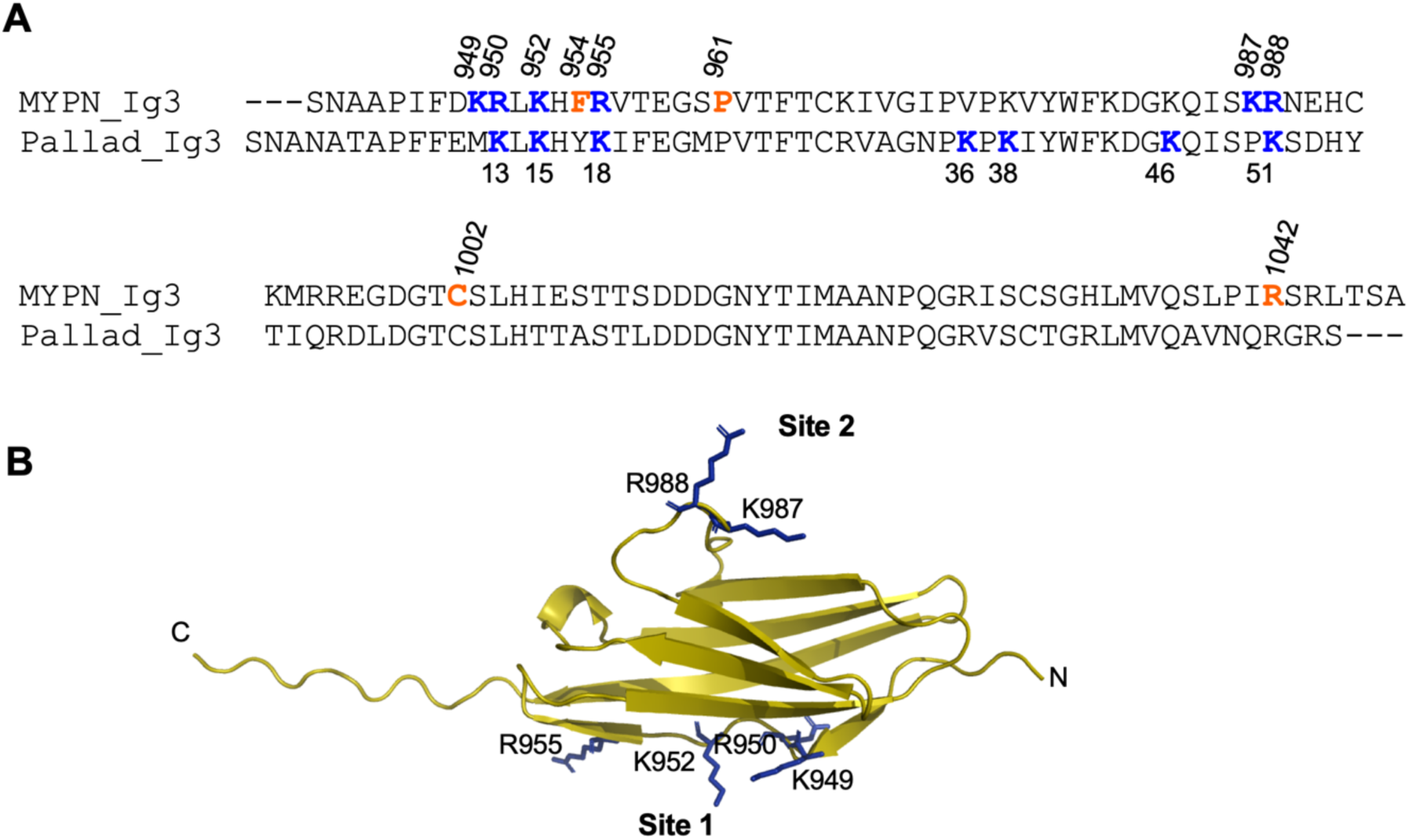
Sequence alignment of MYPN and PALLD Ig3 domains reveals conserved basic patches. A, Alignment shows conserved basic residues (blue). MYPN K952, R955 and R988 correspond to PALLD K15, K18 and K51, respectively. Additionally, PALLD includes other critical residues involved in F-actin interaction (K36, K46, and K38). In MYPN, blue denotes charge-masking mutations; orange denotes a cardiomyopathic mutation (R955 is shared by both mutation types). MYPN Ig3 numbering is based on the full-length human protein, whereas PALLD numbering refers to the isolated mouse Ig3 domain. B, AlphaFold3 predicted structure of the MYPN Ig3, highlighting two basic patches: Site 1 (K949, R950, K952, R955) and Site 2 (K987, R988). Basic side chains are shown as blue sticks.

**Figure 2.**
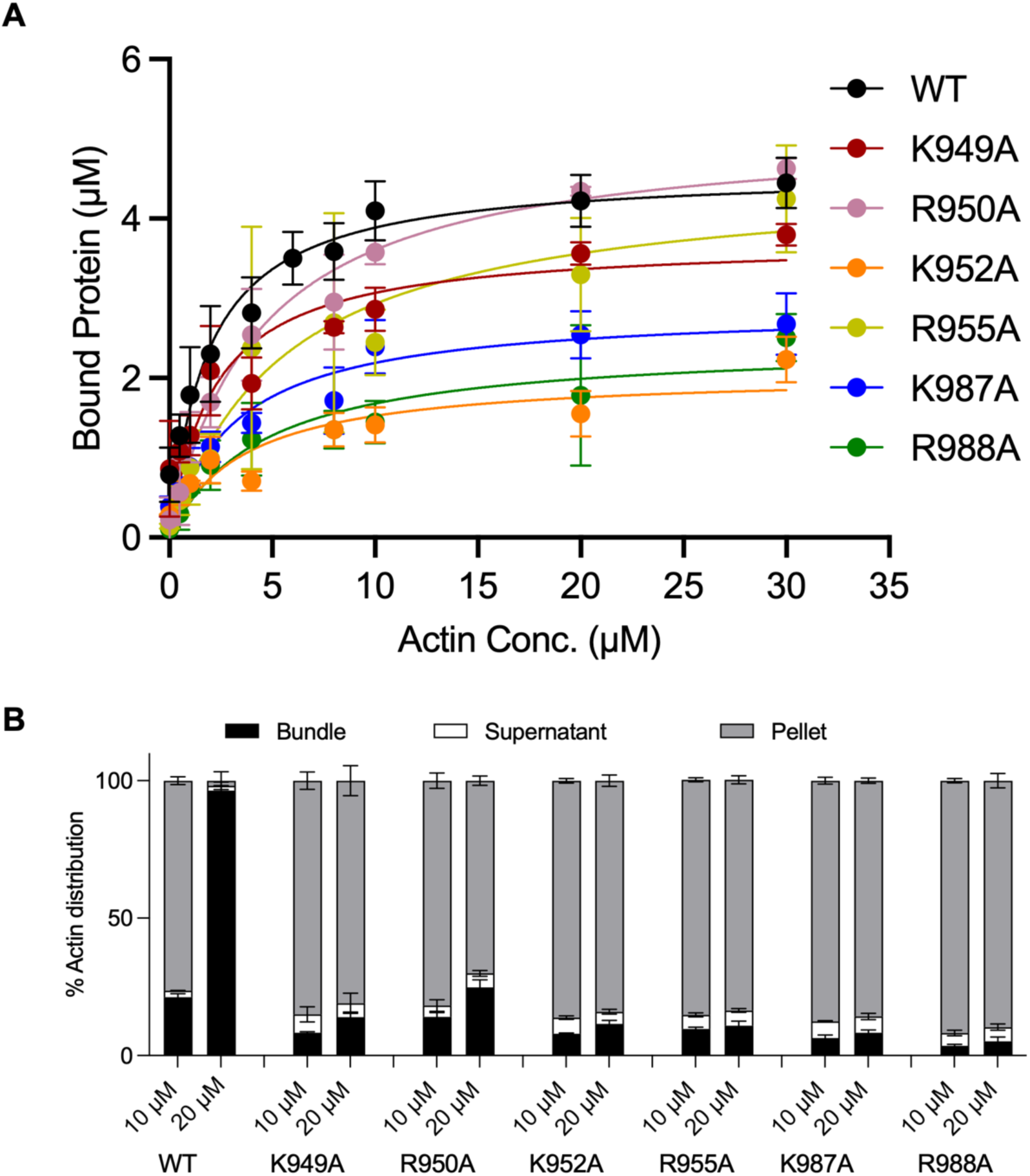
Alanine mutations in conserved basic residues of MYPN Ig3 reduce F-actin binding and impair cross-linking. **A**, Co-sedimentation binding curves where a constant Ig3 concentration (10 µM) was titrated with varying F-actin (0–30 µM). The data were fitted with a hyperbolic curve, assuming specific binding only, to obtain the K_d_ values (Table S1). Data are presented as mean ± SD (n = 3). Mutations reduced apparent K_d_. **B**, Low-speed co-sedimentation bundling assay where constant F-actin concentration (10 µM) was incubated with Ig3 at 1:1 and 1:2 molar ratios. The percentage of pelleted F-actin after low-speed centrifugation represents bundles (black bars). The pellet from high-speed centrifugation represents polymerized F-actin (gray bars). The supernatant contains G-actin (white bars). Data are presented as mean ± SD (n=3). Differences between mutant and WT were considered statistically significant at p < 0.05, as determined by the t-test.

### Cardiomyopathic Mutations Impair Actin Binding and Bundling by MYPN Ig3

The essential contribution of basic residues to F-actin binding and cross-linking, combined with the presence of pathogenic mutations in the Ig3 domain of MYPN^13^ (Figure 3), suggested that these variants might impair MYPN-F-actin interactions and thereby compromise sarcomeric integrity in cardiac muscle. To directly test this hypothesis, we generated site-directed mutants of disease-associated variants, expressed and purified the recombinant proteins, and assessed their actin-binding properties.

**Figure 3.**
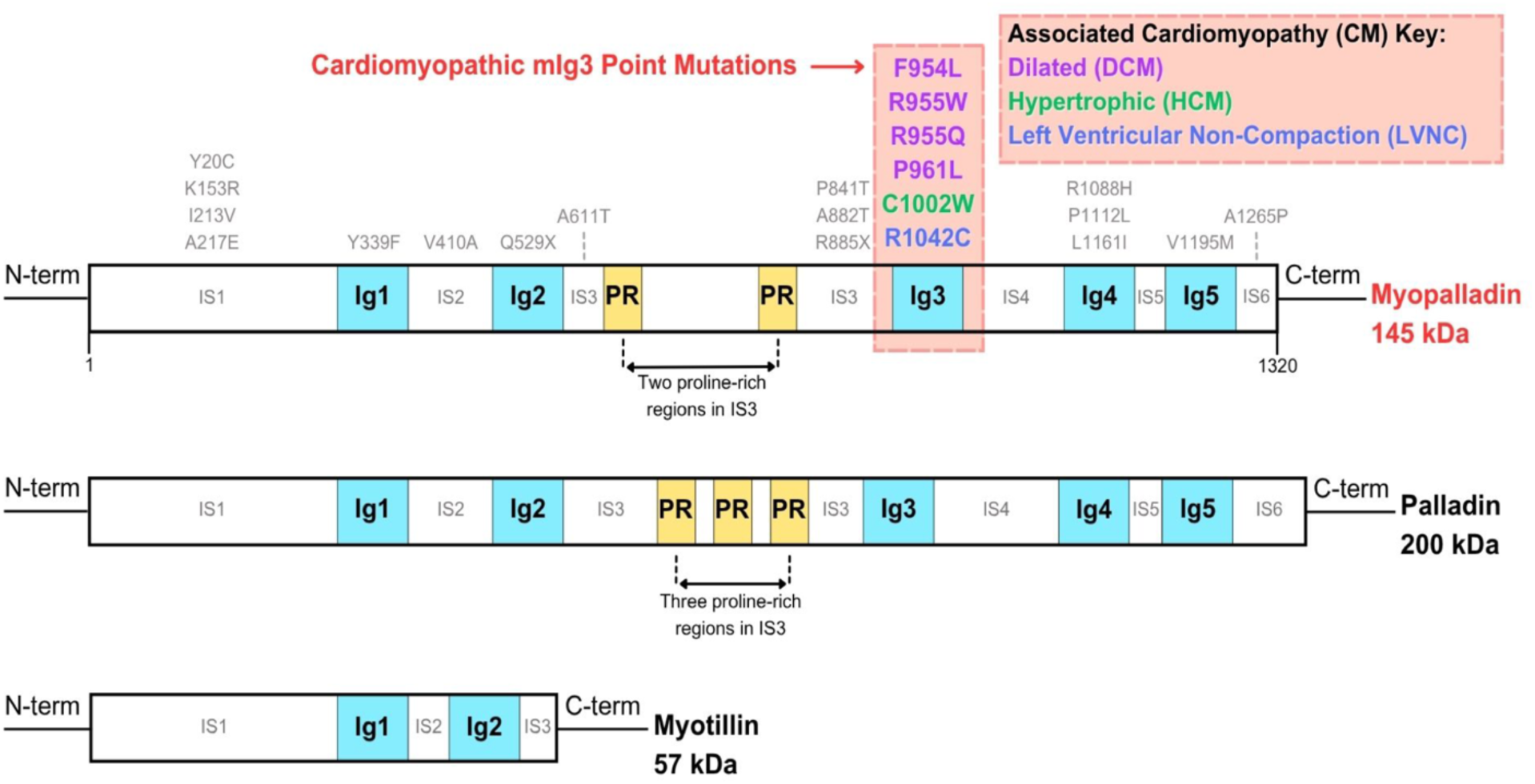
Domain architecture of MYPN and location of CM variants. The domain organization of MYPN is shown along with related proteins (200 kDa isoform 1 of PALLD and myotilin), highlighting Ig domains and polyproline-rich (PR) regions. The locations of a subset of CM variants from the 66 identified CMs for MYPN are indicated. Specific mutations within the Ig3 domain and their associated diseases are detailed in the boxes above MYPN.

Co-sedimentation assays revealed that all CM-associated variants displayed reduced affinity for F-actin compared to WT, as reflected by increased apparent K_d_ values (Figure 4A and Table S1). The P961L mutant could not be quantitatively analyzed due to a high baseline signal in the pellet fraction in the absence of F-actin, consistent with non-specific binding sedimentation or aggregation of the protein. Increasing F-actin concentrations produced only a minimal increase in pelleted P961L, and its markedly reduced F-actin bundling activity further confirmed impaired, non-specific interactions (Figure 4B). Furthermore, circular dichroism (CD) analysis described later indicated that P961L is partially unfolded, consistent with the difficulty in purifying soluble protein and suggesting that misfolding underlies its aggregation propensity.

**Figure 4.**
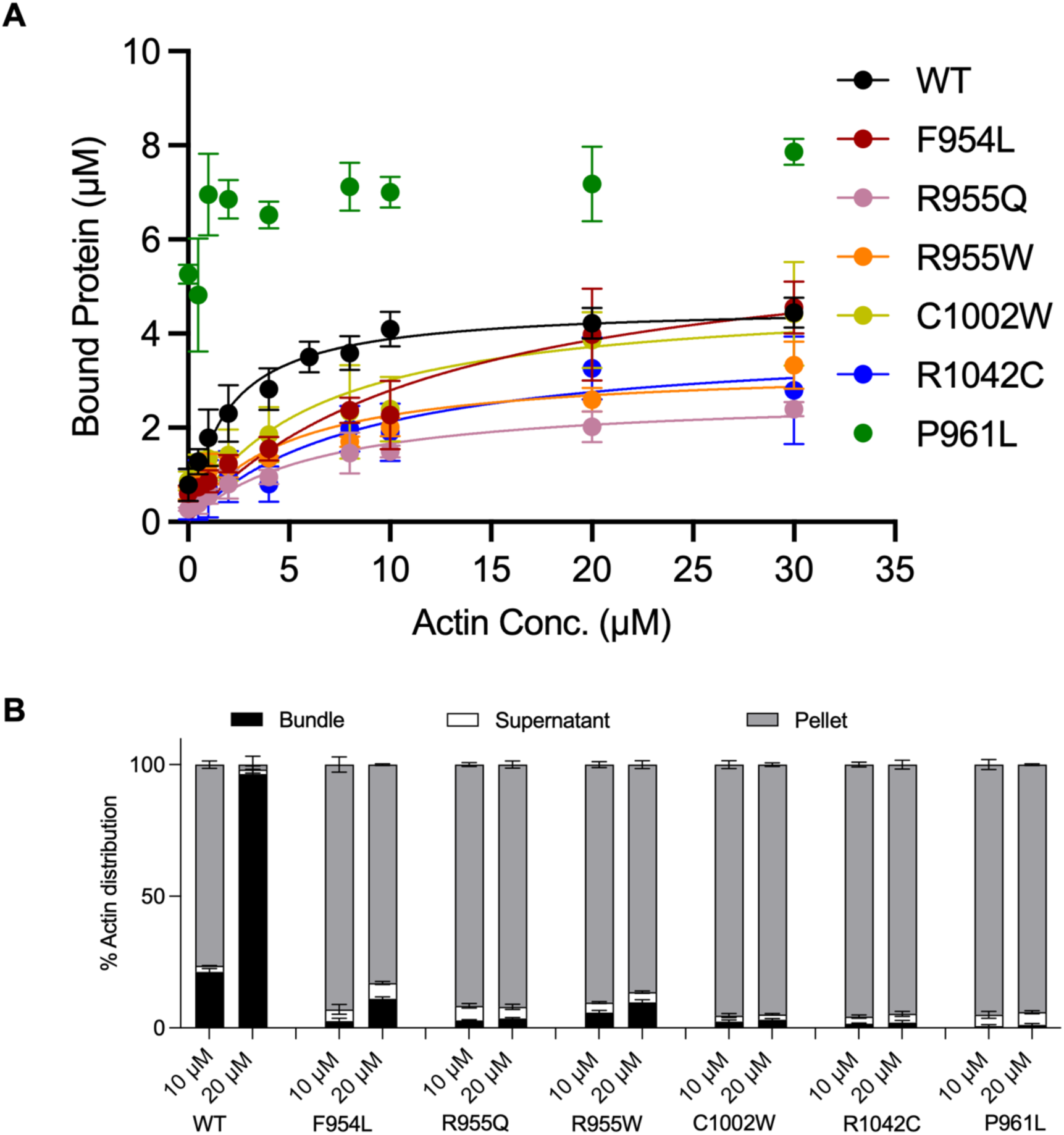
Cardiomyopathy-associated MYPN Ig3 mutants show reduced F-actin binding and dramatic decrease in bundling. **A**, Binding curves generated from co-sedimentation assays where constant Ig3 concentration (10 µM) was titrated with varying F-actin (0–30 µM), showing reduced apparent K_d_. The data were fitted with a hyperbolic curve, assuming specific binding only, to obtain the K_d_ values (Table S1). Data are presented as mean ± SD (n = 3). **B**, Low-speed co-sedimentation bundling assay with constant F-actin concentration (10 µM) mixed with Ig3 at 1:1 and 1:2 molar ratios. The percentage of pelleted F-actin after low-speed centrifugation represents bundles (black bars). The pellet from high-speed centrifugation represents polymerized F-actin (gray bars). The supernatant contains G-actin (white bars). Data are presented as mean ± SD (n=3). Differences between mutant and WT were considered statistically significant at p < 0.05, as determined by the t-test.

Consistent with their weakened actin-binding affinity, all CM-associated variants exhibited a pronounced reduction in actin bundling activity (Figure 4B), reinforcing the conclusion that mutations in the Ig3 domain disrupt MYPN-actin interactions and compromise filament organization.

These results demonstrate that the MYPN Ig3 domain is a functionally critical actin-binding module, and that disease-associated mutations within this region impair MYPN’s capacity to regulate actin filament organization. In particular, the DCM variants F954L, R955Q, and R955W cluster near site 1. The reduced actin-binding function of all three variants highlights this region as a critical actin-interaction interface. Disruption of F-actin engagement at or adjacent to this site provides a mechanistic explanation for Z-line disorganization and compromised sarcomeric architecture observed in cardiomyopathic hearts.

### MYPN Ig3 Promotes Actin Polymerization and Crosslinking During Filament Assembly

Previous studies showed that PALLD Ig3 can crosslink both pre-formed actin filaments and filaments generated during polymerization, with stronger bundling observed when Ig3 is present during filament assembly.^19^ To determine whether MYPN Ig3 exhibits similar behavior, we compared its bundling activity under three conditions: (i) with G-actin (globular actin) in non-polymerizing buffer, (ii) during actin polymerization, and (iii) with pre-formed filaments. As shown in Figure 5, MYPN Ig3 promoted both filament formation and bundling, with bundling efficiency increasing in a concentration-dependent manner across all conditions. Under polymerizing conditions, filaments were formed in actin-only samples, but bundling required Ig3. When Ig3 was added either during polymerization or to pre-formed actin filaments, it effectively bundled F-actin. Unlike the results observed for PALLD Ig3, MYPN Ig3 showed no significant difference in bundling efficiency between co-polymerization and post-polymerization conditions.

**Figure 5.**
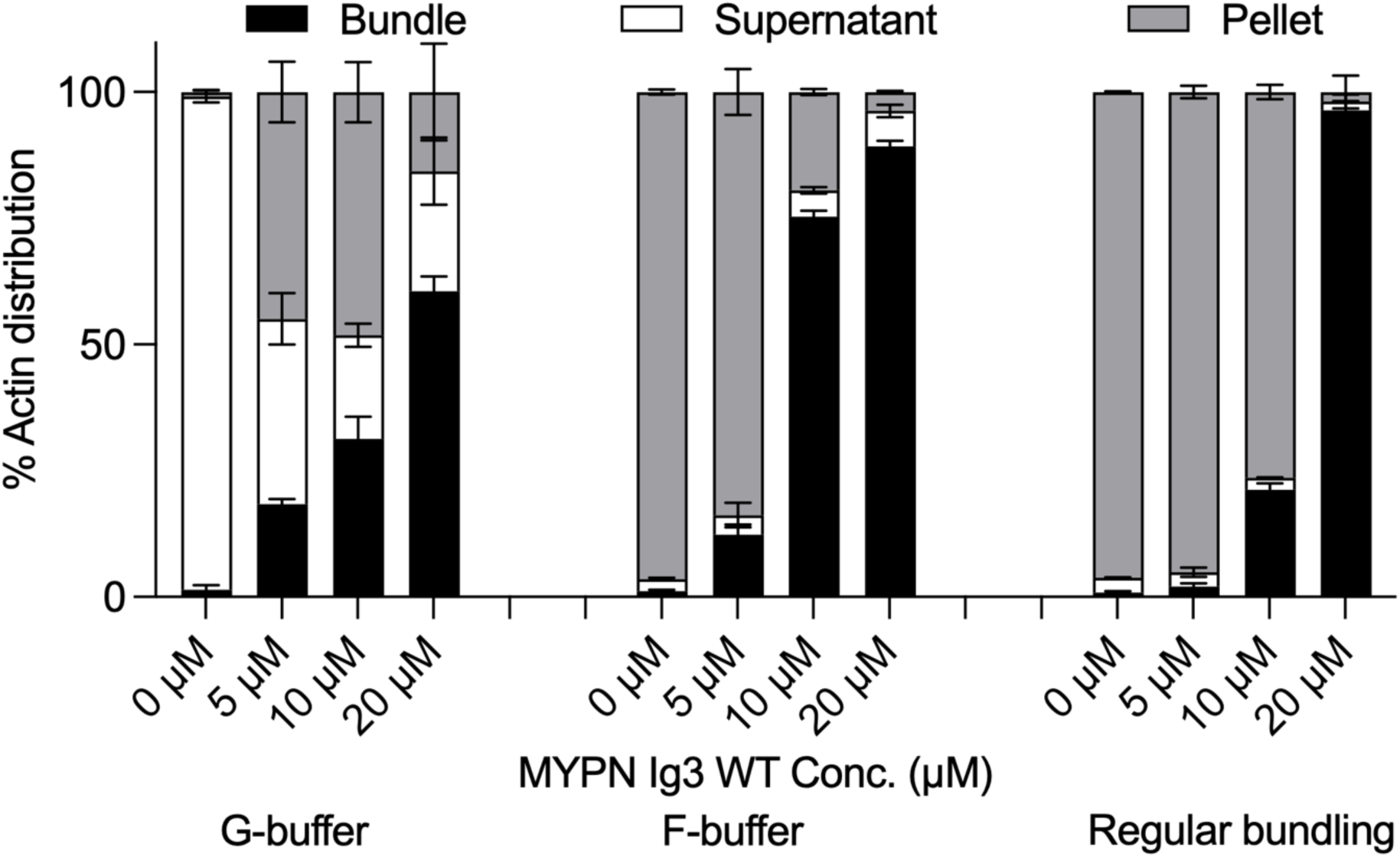
The Ig3 domain polymerizes and bundles G-actin into filaments. G-actin (10 µM) was incubated with increasing concentrations of MYPN Ig3 (0–20 µM) under non-polymerizing (G-buffer) and polymerizing (F-buffer) conditions. For comparison, F-actin (10 µM) was incubated with increasing concentrations of MYPN Ig3 (0–20 µM) under polymerizing conditions (F-buffer). The percentage of actin found in the pellet after low-speed centrifugation is indicated as bundle (black bar), while polymerized actin pelleted at high-speed centrifugation is represented as pellet (gray bar). The G-actin remaining in the soluble fraction is shown as supernatant (white bar). Data are presented as the mean ± SD (n=3). Statistical significance between conditions was assessed using the t-test, with a p-value < 0.05 considered significant.

### Electrostatic Interactions Mediate MYPN Ig3-F-actin Binding

Most actin binding proteins (ABP) depend, to some degree, on electrostatic interactions to engage the negatively charged surface of F-actin. Many of these proteins contain clusters of basic residues within their actin-binding regions. Increasing the ionic strength, by raising the salt concentration, often reduces the affinity for actin due to the disruption of these electrostatic interactions.^20–26^ To determine whether MYPN Ig3-F-actin binding is influenced by ionic strength, we performed co-sedimentation assays across a range of KCl concentrations (25-200 mM).

As shown in Figure 6A, F-actin binding was highest at 100 mM KCl, which is the standard concentration used in prior assays, and significantly reduced at 50 mM and 200 mM (p< 0.05). Binding at 25 mM KCl was similar to that at 100 mM (not significantly different at *p* < 0.05), suggesting that both low and physiological ionic strengths support interaction, but high salt disrupts it. These results indicate that electrostatic interactions contribute to MYPN Ig3 F-actin binding, and that this interaction is partially weakened at elevated ionic strength, similar to PALLD Ig3.^26^ Interestingly, MYPN Ig3 exhibited maximal binding at physiological ionic strength (100 mM KCl), unlike PALLD Ig3, which binds most strongly at low salt. This suggests that MYPN may have evolved distinct electrostatic tuning for stable interaction with actin in the cardiac cytoskeletal environment.

**Figure 6.**
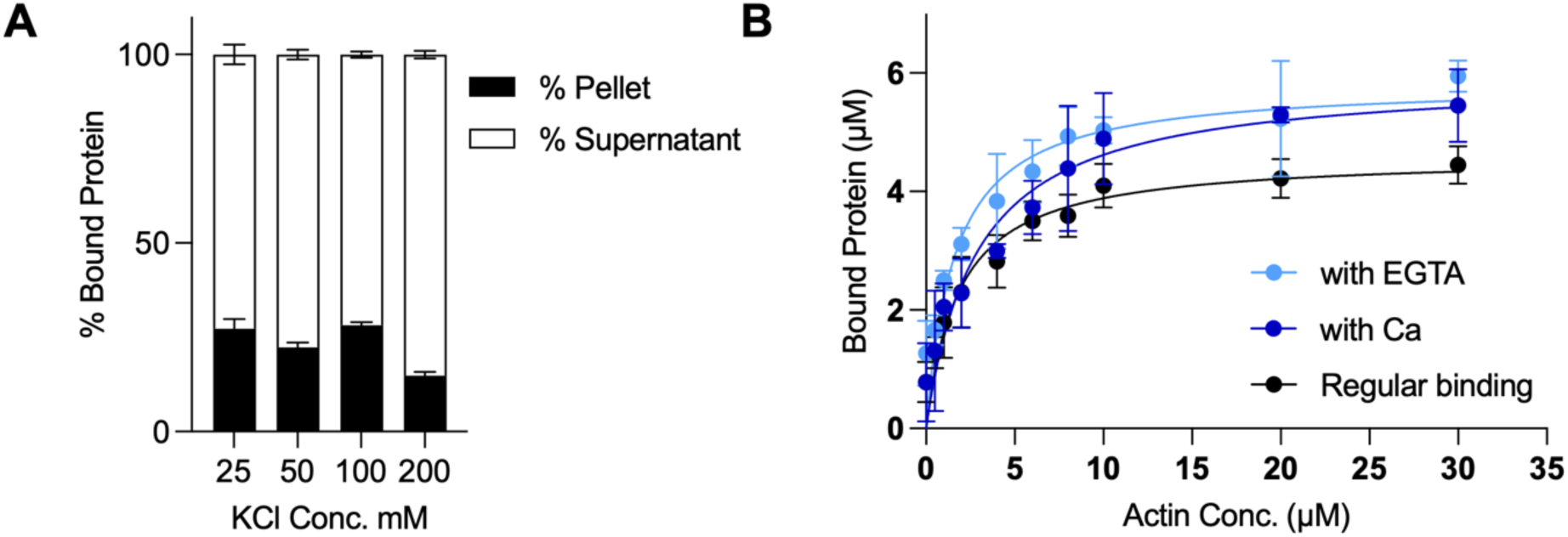
MYPN Ig3-F-actin interaction is electrostatically driven but not modulated by Ca^2+^. A, Co-sedimentation of 20 µM Ig3 with 5 µM F-actin at increasing KCl concentrations (25, 50, 100, and 200 mM). The percentage of Ig3 bound to F-actin (pellet fraction) is shown in black, while the unbound Ig3 (supernatant fraction) is shown in gray. B, Binding curves of Ig3 (10 µM) titrated with F-actin (0–30 µM) with Ca^2+^ or EGTA. Apparent K_d_ values: 2.86 ± 1.15 µM (+Ca^2+^), 1.61 ± 0.60 µM (-Ca^2+^, pCa=10), and 1.80 ± 0.60 µM (standard conditions). Data are presented as the mean ± SD (n = 3). Statistical significance between conditions was assessed using the t-test, with a p-value < 0.05 considered significant.

### MYPN Ig3 Binding to Actin is Independent of Calcium

Upon muscle cell activation, sarcoplasmic Ca^2+^ concentrations rise from a resting concentration of ∼100 nM to ∼1 µM.^27^ Previous studies have shown that the N2A domain of titin, a sarcomeric ABP, exhibits Ca^2+^-dependent binding to F-actin.^28^ To determine whether the MYPN Ig3 domain is similarly regulated by Ca^2+^, we performed co-sedimentation assays under Ca^2+^-rich (pCa = 4) and Ca^2+^-free (EGTA, pCa = 10) conditions. As shown in Figure 6B, the apparent K_d_ of MYPN Ig3 for F-actin was not significantly affected by Ca^2+^. The K_d_ was 2.86 ± 1.15 µM in the presence of Ca^2+^ and 1.61 ± 0.60 µM in the absence of Ca^2+^, both comparable to the baseline K_d_ under standard assay conditions (1.84 ± 0.63; *p* < 0.05).

These results indicate that MYPN Ig3-actin binding is Ca^2+^-independent, suggesting a constitutive role in actin cytoskeleton maintenance. This stable interaction may help preserve Z-line integrity during rapid Ca^2+^ fluctuations that occur with muscle contraction. Such stability could also protect binding interfaces for other regulatory proteins, such as titin, that respond dynamically to calcium signalling.

### Most Cardiomyopathy-linked and Charge-Neutralizing Mutations Preserve the Ig Fold of MYPN

To determine whether any of the point mutations affect the secondary structure of the MYPN Ig3 domain, we performed far-UV CD spectroscopy. The CD spectrum of WT Ig3 exhibited a maximum near 205 nm and a minimum around 215 nm, consistent with the signature of antiparallel β-sheets, which typically exhibit a peak at 195 nm and a trough at 218 nm.^29^ These results confirm the β-sheet-rich architecture of the Ig3 domain in MYPN. Notably, the spectral features of MYPN Ig3 are slightly shifted from canonical β-sheet signatures, likely due to the β-sandwich topology of this domain, in which two opposing β-sheets are stacked together.^30,31^ This geometry, along with strand twisting,^32^ can influence CD signal positions. The CD spectra of all MYPN Ig3 mutants closely overlap with WT (Figure S2), indicating that secondary structure remains intact. Minor deviations were observed for R950A and C1002W but did not reflect any major structural disruption. These data suggest that neither disease-associated nor charge-neutralizing mutations disrupt the β-sheet architecture of the MYPN Ig3 domain. Protein secondary structure content of WT Ig3 domain was estimated with CDSSTR to contain ∼23% regular β-sheet (strand-1), ∼14% distorted β-sheet (strand-2), and negligible α-helical content (<5%), consistent with β-sandwich topology^30,31^. CDSSTR analysis of R950A and F954L mutants showed no significant deviations from the WT.^33,34^ K2D estimates for all variants were consistent including the WT, yielded ∼10% α-helix, ∼39-47% β-sheet, and ∼44-49% random coil (Table S3). Collectively, these results indicate that neither CM-related nor charge-modifying mutations disrupt the secondary structure or hydrophobic core packing of the Ig3 domain, preserving the integrity of the Ig fold in all tested variants.

Unlike the WT and other variants, the P961L mutant lacked the characteristic β-sandwich spectrum (Fig. S2B). Its secondary structure profile was altered relative to WT, showing reduced b-sheet content (38%) and increased random coil (50%), while a-helix content remained similar (12%). These results suggest that P961L mutation disrupts the ability of the domain to adopt the canonical Ig fold.

### Mutations Modulate Thermal Stability Without Disrupting Folding

Although CD spectroscopy confirmed that most disease-associated and charge-neutralizing mutations do not alter the secondary structure of MYPN Ig3, subtle differences in domain stability could still affect function or increase susceptibility to misfolding under physiological stress. Notably, the CM mutation P961L did not adopt a stable folded structure. To assess these effects, we measured the thermal stability of WT and mutant Ig3 domains using CD thermal denaturation. WT MYPN Ig3 unfolded cooperatively with a thermal denaturation midpoint (T_M_) of 48.50 ± 0.85 °C (Figure 7). Most mutants showed altered thermal stability. Variants K987A and F954L exhibited modest but significant (p < 0.05) and reproducible increases in T_M_ (50.48 ± 0.20 and 50.11 ± 0.21 °C, respectively), consistent with enhanced stability through modified intramolecular interactions. In contrast, C1002W and R950A displayed significantly reduced thermal stability (p < 0.05; T_M_ = 45.10 ± 0.33°C and 42.38 ± 0.34 °C) and less cooperative unfolding transitions, indicative of local destabilization or increased conformational flexibility. Interestingly, the CM-associated variant R955W, located near the actin-binding surface, exhibited an intermediate state in its thermal profile before full denaturation. However, its T_M_ value (48.05 ± 0.45 °C), was not significantly different from that of the WT. In contrast, the T_M_ of P961L could not be determined, as its CD spectrum resembled that of an unfolded protein, and the melting curve was flat, showing no change between folded and unfolded states. No significant changes in thermal stability were observed for the remaining mutants (Table S3).

**Figure 7.**
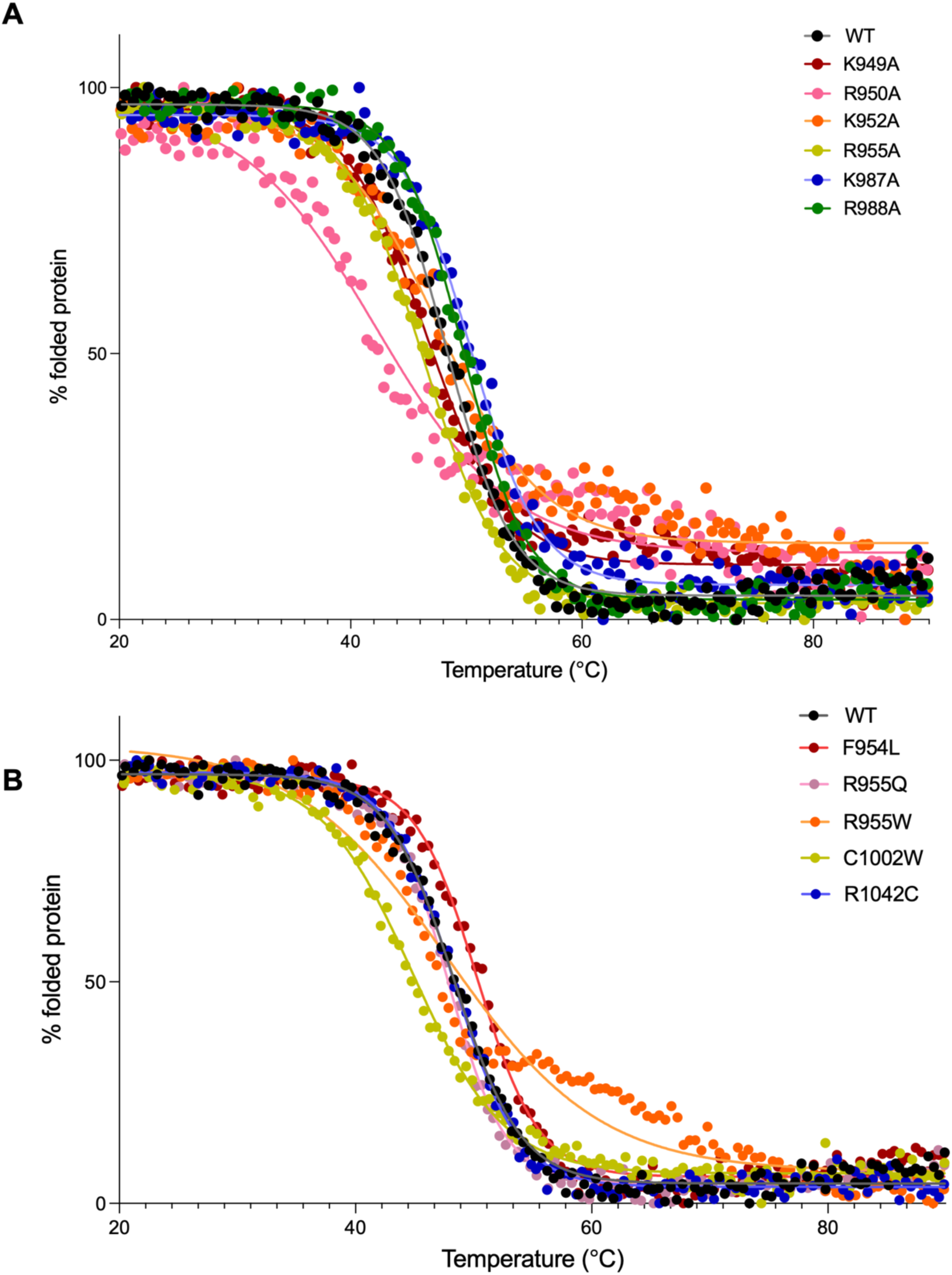
Mutations alter MYPN Ig3 thermal stability. Thermal denaturation of WT and mutant Ig3 was monitored by CD at 205 nm. Data were globally fit to a two-state unfolding model to determine the T_M_ and Hill coefficient. **A**, Unfolding curves for alanine mutations and **B**, CM mutations, compared with WT.

Overall, these results indicate that the MYPN Ig3 domain is structurally robust, tolerating many substitutions without major disruption, although P961L is a clear exception. However, specific mutations subtly modulate thermodynamic stability, which may alter folding, interaction dynamics, or stability *in vivo*. Similar effects have been observed in other Ig-like domains, such as titin I21, where single-residue substitutions (e.g., C3575S) decreased stability without altering the overall fold.^35^ By integrating structural and thermodynamic analyses, we demonstrate that the MYPN Ig3 domain tolerates many mutations without major structural disruption, yet select variants fine-tune its stability in ways that may contribute to cardiomyopathy pathogenesis.

### MYPN Ig3 Domain Exists as a Monomer in Solution

To determine the oligomeric state of the MYPN Ig3 domain in solution, we performed sedimentation equilibrium analytical ultracentrifugation (SE-AUC), a quantitative method for measuring native molecular mass in solution. Nonlinear least-squares fitting of the concentration gradients to a self-association model revealed no evidence of higher-order species (Figure S3). Across multiple protein concentrations and rotor speeds, the measured molecular mass was 11.7 kDa, closely matching the predicted monomeric mass of 12 kDa, and showed no concentration-dependent increase in molecular weight. These results indicate that MYPN Ig3 domain exists as a stable monomer in solution under our experimental conditions. This is consistent with previous findings for the homologous PALLD Ig3 domain, which also functions as a monomer despite its actin-bundling capability.^15,26,36^ This suggests that multivalent F-actin interactions are achieved through the structural topology or surface chemistry of individual Ig3 domains, rather than via domain organization.

In contrast, other Ig-like domains, particularly in structural proteins such as filamin and titin, can mediate dimerization or form elongated multimers. For example, certain filamin Ig domains participate in homodimeric or heterodimeric associations that are essential for actin cross-linking and mechanical resistance.^37^ Similarly, titin’s Ig and FnIII domains can form supramolecular assemblies that contribute to sarcomeric elasticity.^38^ The fact that MYPN Ig3 and its PALLD homolog, retain actin-bundling activity as monomers highlights a distinct molecular strategy involving the use of highly specialized binding interfaces rather than multimerization to achieve filament crosslinking. This may reflect unique functional constraints at the Z-line, where tight spatial packing of sarcomeric proteins requires precise, monomeric interactions.

### MYPN Function in Drosophila Muscle Tissues

To investigate the impact of MYPN mutations *in vivo*, we generated humanized *Drosophila* models to analyze MYPN protein localization in larval heart and muscle tissues. The *Drosophila* genome does not encode for MYPN, so we expressed wild-type and mutant versions in transgenic fly lines utilizing the GAL4-UAS binary expression system.^39^ The *Mef2* promoter drives expression of the transactivator GAL4 protein in heart and skeletal muscle throughout the *Drosophila* life cycle.^40^ GAL4 binds to upstream activating sequences (UAS) to induce expression of MYPN proteins.

Immunostaining of the larvae heart tube revealed localization of human MYPN to the Z-disc of cardiac tissue using our GAL4-mediated expression system (*Mef2*-GAL4>UAS-*MYPN WT* (Figure 8A). This result nicely validates the *Drosophila* musculature as an evolutionary conserved model to study mammalian muscle structure and function as human MYPN is also located at the Z-disc.^41^ The *Drosophila* larval body wall muscles are larger than the heart tube and the organization of sarcomeres in series within each muscle cell are ideal for studying sarcomeric protein localization. As expected due to the lack of MYPN in *Drosophila*, there was no MYPN signal in *w^1118^* control muscles (Figure 8B). Expression of human WT MYPN is located at the Z-disc (Figure 8B, open white triangle) in larval muscles. To verify our immunostaining results, we also generated transgenic flies containing GFP-tagged versions of MYPN and found that MYPN-GFP showed the same Z-disc localization as the untagged protein (Figure S2B). Importantly, the overall pattern of larval muscles was not altered upon expression of MYPN or MYPN mutant proteins appended with GFP (Figure S2A).

**Figure 8.**
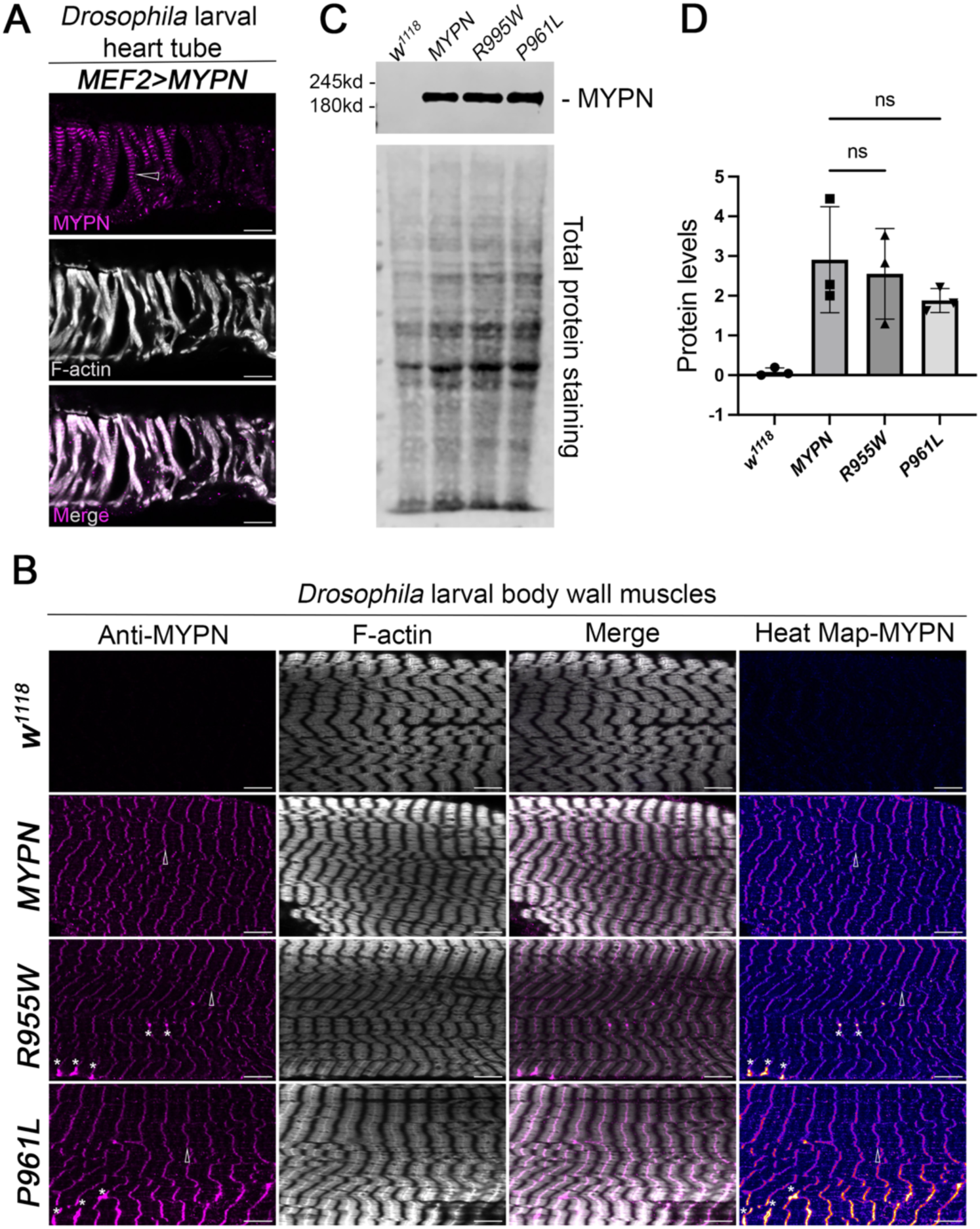
MYPN is located at the Z-disc and Ig domain mutations cause aberrant MYPN accumulation. **A**, Immunostaining of *Drosophila* larval heart tube expressing *Mef2>MYPN WT*. MYPN protein is localized to the Z-disc (magenta, open white triangle). F-actin is shown in grey. Scale bar: 10mM. **B**, L3 larval muscles stained with phalloidin to visualize F-actin or with an antibody against MYPN. F-actin and MYPN co-localize at the Z-disc (open white triangle), in *Mef2>MYPN WT* muscles. *MYPN R955W* and *MYPN P961L* Ig domain mutations show abnormal accumulation of MYPN (white asterisk). Scale bar: 10mM. **C**, Western blot showing the expression levels of MYPN^62^ in the indicated genotypes. Total protein staining serves as a loading control. One-way ANOVA. n = 3. **D**, Quantification of MYPN expression levels.

In *MYPN R955W* and *P961L* mutants, we discovered that MYPN was largely present at its expected location (Figure 8B, open white triangle) but also accumulated abnormally in clusters away from the Z-disc or appeared as widened Z-discs (Figure 8B, asterisks). Similar phenotypes were observed for GFP-tagged MYPN mutants, where R955W-GFP signal extended into the thin filament (Figure S2B, open white arrow) and P961L-GFP resulted in larger aggregates (Figure S2B, asterisks). Overexpression of all MYPN-GFP proteins resulted in substantial accumulation near the nuclei (Figure S2C). Western blotting was used to confirm expression levels of all MYPN proteins and confirmed that the differences in subcellular localization were not due to differences in expression levels (Figures 8C-D; S3D-E).

## DISCUSSION

The identification of sixty-six MYPN missense variants associated with DCM, HCM, RCM, and LVNC cardiomyopathies^13,42^, combined with observed myofibrillar abnormalities in patients^14^ and MYPN’s interactions with key sarcomeric structural components, collectively underscores its role in maintaining sarcomere integrity and normal cardiac function.^4,9^ Since the underlying disease mechanisms remain poorly understood and the functional role of MYPN in the heart has remained elusive, the development of new therapies for cardiomyopathies has been hampered. Previous studies have primarily focused on MYPN’s role in signaling pathways and nuclear shuttling disruption.^8,10^ These investigations examined specific MYPN mutations (Y20C and Q529X) located in the region that interacts with cardiac ankyrin repeat protein (CARP/Ankrd1), a stress-inducible transcriptional co-factor that suppresses genes involved in heart failure and hypertrophy development.^8,10^ Our study focuses on disease-associated mutations within MYPN’s actin-binding region, a domain that directly involves thin filament binding and assembly. To better understand MYPN’s role in cardiac function, we characterized how the Ig3 domain modulates actin dynamics and determined the functional alterations caused by CM mutations.

Ig domains are evolutionarily conserved protein-protein interaction modules present in several cytoskeleton-associated proteins^31^, including MYPN, myomesin,^43^ titin,^44^ MyBP-C,^45^ MyBP-H,^45^ filamin C,^46^ obscurin,^47^ PALLD,^15^ and myotilin.^48^ Although these domains share a similar overall structure, they engage with a wide range of binding partners, from actin and myosin to nuclear components, prompting the question of what structural features underlie their specificity. Several known actin-binding motifs, such as calponin homology domain, Wiskott-Aldrich syndrome protein homology domain-2 motifs, gelsolin homology, actin-depolymerizing factor homology, and the lysine-rich actin-binding domain 3 (ABD3) motif, rely on clusters of positively charged residues for actin interaction.^15,49–52^ PALLD, MYPN, and myotilin constitute a closely related subfamily of Ig-domain containing proteins, and in both PALLD and myotilin, F-actin binding is mediated by conserved basic clusters. Specifically, myotilin relies on residues K354, K358, and K359,^48^ while PALLD contains lysine-rich sequences (K13, K18, K36, K46 and K51), where substitution of these residues abolishes F-actin binding.^15,16^

To elucidate the molecular basis of MYPN’s regulation of actin dynamics, we examined the role of positively charged residues in the Ig3 domain’s actin-binding region. Charge-neutralizing mutational analysis revealed that two surface-exposed basic patches on opposing faces of the Ig3 domain are essential for actin binding. We identified two primary actin-binding sites: one comprising K949, R950, K952, and R955, and another containing K987 and R988. Mutations in either site significantly reduced F-actin binding affinity, but no single mutation completely abolished the interaction, suggesting additional residues also contribute to binding. Thus, MYPN, like its related family members, relies on clusters of surface-exposed basic residues to mediate F-actin binding.^15,48^

In cells, actin filaments assemble into diverse higher-order architectures, and actin cross-linking proteins can regulate this structural complexity.^53^ Cross-linking generally requires either two actin-binding sites within a single protein (e.g., PALLD, fascin, fimbrin, and espin) or dimerization of monomeric proteins that contain only one binding site (e.g., myotilin).^53,54^ Our mutational data show that neutralizing either basic patch significantly impaired actin cross-linking, mirroring effects observed with analogous mutations in PALLD’s Ig3 domain.^15^ Our data highlight the indispensable role of these residues in actin bundle formation. Supporting this, our AUC analysis demonstrated that the isolated Ig3 domain exists as a monomer in solution. We therefore propose that the two basic clusters on opposite faces of the Ig3 domain permit a single MYPN molecule to simultaneously engage and bundle two actin filaments.

Our *in vitro* data on CM causing mutations demonstrate that all variants impair F-actin binding and have a significant effect on actin bundling. This aligns with data from the muscle-specific α-actinin-2 ABD, in which HCM-associated mutations (A119T and G111V) also reduce F-actin affinity while preserving secondary structure.^55^ Among the DCM-causing mutants, P961L has the most dramatic effect, abrogating specific actin interactions. This trend of impaired actin-binding extends to other DCM-associated mutants (F954L, R955Q, and R955W), an HCM-associated mutant (C1002W), and an LVNC-associated mutant (R1042C), suggesting a common pathological mechanism for cardiomyopathies found in cardiomyopathies.

Expression of human WT MYPN in *Drosophila* larval muscles demonstrated proper Z-line localization, despite the absence of endogenous *MYPN* in the fly genome. Immunostaining and GFP-tagged constructs revealed that the R955W and P961L mutant proteins accumulated into clusters, incorporating improperly into Z-discs. Notably, GFP-P961L exhibited prominent protein aggregates. Anti-MYPN staining further demonstrated widened Z-discs and, in the case of GFP-R955W, partial diffusion of MYPN into thin filaments. Consistent with these findings, the biopsies from DCM patients, the R955W showed normal staining patterns, however P961L exhibited profound pathological effects,^14^ disrupting MYPN’s interactions with α-actinin and leading to abnormal protein distribution and altered cytoskeletal organization with severely compromised sarcomere structure.^14^ These findings demonstrate how CM mutations disrupt MYPN’s interactions with both actin and other sarcomeric proteins, leading to Z-disc structural defects representing a likely mechanism underlying their pathological effects.

CD analysis revealed that most mutants, except P961L, retained secondary structure content similar to WT, maintaining the Ig-fold β-sandwich. However, melting temperature measurements indicated that some mutations altered thermal stability without disrupting the overall fold. In contrast, the CD spectrum of the P961L mutant resembled that of an unfolded protein, indicating that this mutation abolishes its characteristic Ig-fold. This mutation likely destabilizes the hydrophobic core of the domain, suggesting that the proline residue at 961 is critical for maintaining the correct fold of the Ig3 domain. Along with the tissue staining patterns in both *Drosophila* and human samples, these findings suggest that the P961 residue is critical for maintaining the structural integrity of full-length MYPN, which in turn is essential for its actin-binding function.

Although the MYPN Ig3 domain has been shown to inhibit actin polymerization in a concentration-dependent manner under polymerizing conditions,^12^ its role in *de novo* assembly of actin monomers into bundled structures has not yet been characterized. Our experiments with G-actin co-polymerization revealed that MYPN Ig3 can induce actin polymerization even under non-polymerizing conditions, subsequently promoting filament bundling. Notably, we observed no significant difference in bundling efficiency between Ig3 co-polymerized with G-actin and its interaction with pre-polymerized or mature actin filaments. This behavior contrasts with that of PALLD Ig3, which generates more extensive actin bundling when co-polymerized with G-actin than with mature filaments.^19^ These results suggest that, unlike PALLD Ig3, MYPN Ig3 facilitates actin bundling independently of the polymerization state of the filaments, indicating a distinct mechanism of actin network regulation.

The electrostatic model of F-actin binding has been proposed for many ABPs, as these typically contain clusters of positively charged residues on their solvent-exposed surfaces, while F-actin itself possesses strongly anionic subdomains.^15,56^ Furthermore, the polyelectrolyte nature of F-actin is known to be stabilized by long-range electrostatic interactions.^57,58^ To determine whether MYPN–F-actin binding is electrostatically driven, we examined the salt sensitivity of the MYPN Ig3–F-actin interaction. We found that MYPN-actin interactions are strongest at physiological salt concentrations (100 mM KCl) and lower salt (25 mM KCl) but are significantly reduced at higher salt concentrations (150 mM KCl). This salt-dependent reduction is consistent with electrostatic contributions. Interestingly, MYPN Ig3 behaves differently from closely related proteins PALLD, myotilin, and coronin 1B. Its binding at 100 mM KCl remains just as strong as binding at 25 mM KCl, whereas F-actin binding by these other ABPs consistently decreases with increasing salt concentration.^26,48,59^ Nevertheless, the reduced co-sedimentation of MYPN with F-actin at elevated salt concentrations supports a role for electrostatic interactions in MYPN–F-actin binding.

The interaction between actin and certain ABPs, such as including filamin A, titin (T2 fragment and N2A region), and non-muscle α-actinins, is known to be regulated by calcium.^28,60,61^ Given that MYPN ablation does not alter myofilament calcium sensitivity,^12^ we examined whether the presence of MYPN influences F-actin binding in a calcium-dependent manner, as seen with other ABPs. Our experiments demonstrate that calcium does not modulate the binding between MYPN’s Ig3 domain and F-actin. Thus, the calcium-independent interaction between the Ig3 domain and F-actin could further explain the lack of effect on myofilament calcium sensitivity upon its ablation.

In summary, this study provides a potential mechanistic link between MYPN mutations and cardiomyopathy, revealing how disruption of MYPN’s actin-binding function drives aberrant accumulation of mutant proteins at the Z-disc and compromises sarcomere integrity. Importantly, we uncover that the Ig3 domain harbors two strategically positioned, solvent-exposed actin-binding sites, underscoring its pivotal role in orchestrating actin dynamics. Together, these findings position the Ig3 domain as a critical nexus in cardiac cytoskeletal regulation and a potential focal point for understanding and targeting MYPN-associated CMs.

## Nonstandard Abbreviations and Acronyms

ABD: actin binding domain
ABP: actin binding protein
CARP: cardiac ankyrin repeat protein
CD: circular dichroism
CM: cardiomyopathy
DCM: dilated cardiomyopathy
F-actin: filamentous actin
G-actin: globular actin
GFP: green fluorescent protein
HCM: hypertrophic cardiomyopathy
Ig: immunoglobulin
LVNC: left ventricular noncompaction cardiomyopathy
MBP: maltose binding protein
ME: molecular ellipticity
MYPN: myopalladin
PALLD: palladin
RCM: restrictive cardiomyopathy
SDS-PAGE: sodium dodecyl sulfate poly acrylamide gel electrophoresis
TEV: tobacco etch virus
WT: wild-type

## ACKNOWLEDGMENTS

The authors thank Dr. Jim Bann, Wichita State University, for assistance with CD data analysis. The MYPN antibody was kindly provided by Marie-Louise Bang, Institute of Biomedical and Genetic Research – National Research Council.

## MATERIAL

Supplemental Methods

Tables S1-S3

Figure S1-S3

Major Resources Table

---

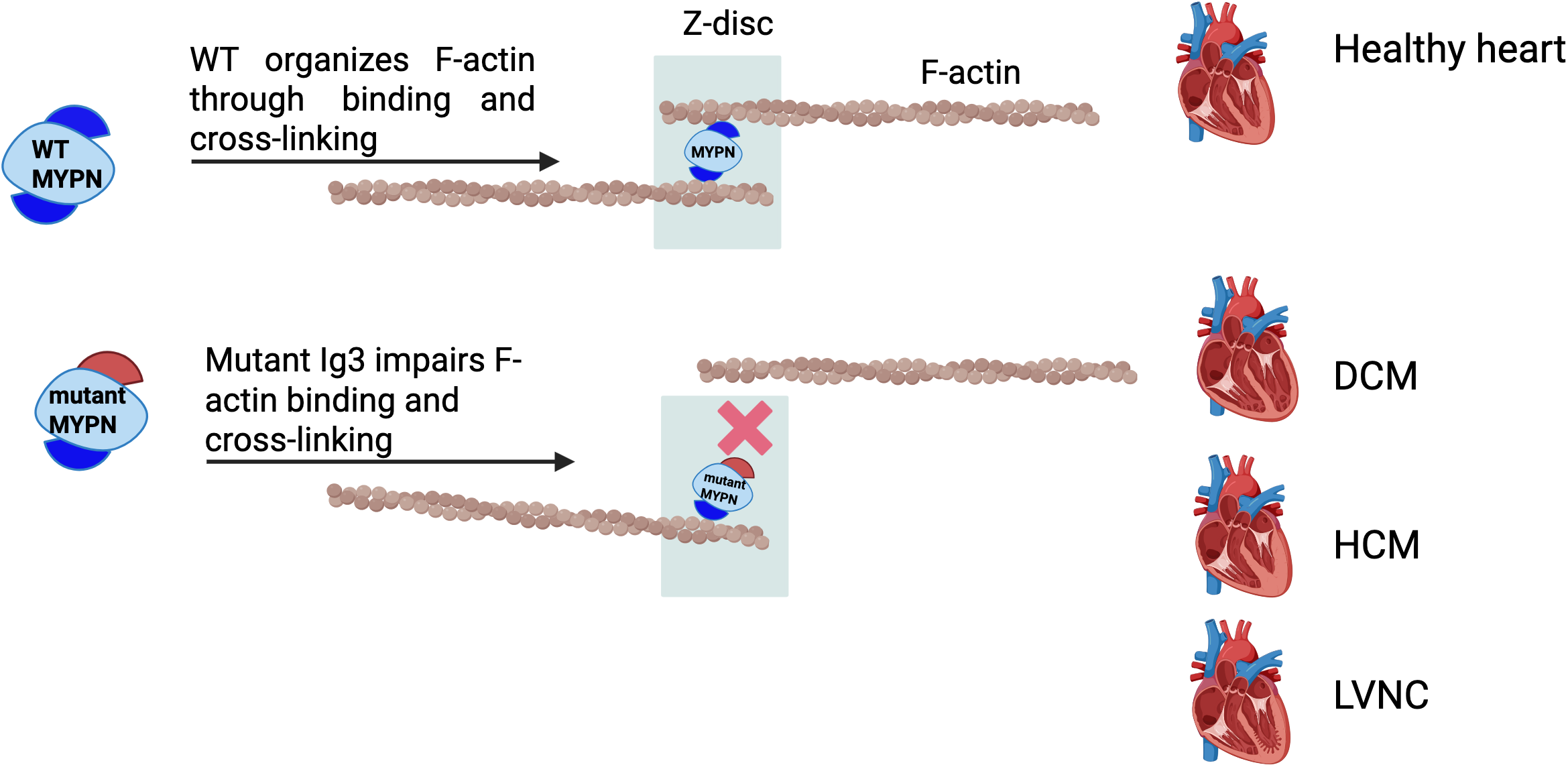

## REFERENCES

1. Towbin J, Bowles N. The failing heart. Nature. 2002;415:227–233.

2. Maron BJ, Towbin JA, Thiene G, Antzelevitch C, Corrado D, Arnett D, Moss AJ, Seidman CE, Young JB. Contemporary definitions and classification of the cardiomyopathies: an American Heart Association scientific statement from the council on clinical cardiology, heart failure and transplantation committee; quality of care and outcomes research and functional genomics and translational biology interdisciplinary working groups; and council on epidemiology and prevention. Circulation. 2006;113:1807–1816.

3. Elliott P, Andersson B, Arbustini E, Bilinska Z, Cecchi F, Charron P, Dubourg O, Kühl U, Maisch B, McKenna WJ. Classification of the cardiomyopathies: a position statement from the European Society Of Cardiology Working Group on Myocardial and Pericardial Diseases. European heart journal. 2008;29:270–276.

4. Bang M-L, Bogomolovas J, Chen J. Understanding the molecular basis of cardiomyopathy. American Journal of Physiology-Heart and Circulatory Physiology. 2022;322:H181–H233.

5. Seidman J, Seidman C. The genetic basis for cardiomyopathy: from mutation identification to mechanistic paradigms. Cell. 2001;104:557–567.

6. Martin SS, Aday AW, Almarzooq ZI, Anderson CA, Arora P, Avery CL, Baker-Smith CM, Barone Gibbs B, Beaton AZ, Boehme AK. 2024 heart disease and stroke statistics: a report of US and global data from the American Heart Association. Circulation. 2024;149:e347–e913.

7. Statistics NCfH, Statistics NCfH. Multiple Cause of Death 2018–2022 on CDC WONDER Database. *National Center for Health Statistics: Hyattsville, MD*, USA. 2023.

8. Purevjav E, Arimura T, Augustin S, Huby A-C, Takagi K, Nunoda S, Kearney DL, Taylor MD, Terasaki F, Bos JM. Molecular basis for clinical heterogeneity in inherited cardiomyopathies due to myopalladin mutations. Human molecular genetics. 2012;21:2039–2053.

9. Filomena MC, Yamamoto DL, Carullo P, Medvedev R, Ghisleni A, Piroddi N, Scellini B, Crispino R, D’Autilia F, Zhang J. Myopalladin knockout mice develop cardiac dilation and show a maladaptive response to mechanical pressure overload. Elife. 2021;10:e58313.

10. Huby A-C, Mendsaikhan U, Takagi K, Martherus R, Wansapura J, Gong N, Osinska H, James JF, Kramer K, Saito K. Disturbance in Z-disk mechanosensitive proteins induced by a persistent mutant myopalladin causes familial restrictive cardiomyopathy. Journal of the American College of Cardiology. 2014;64:2765–2776.

11. Bang M-L, Mudry RE, McElhinny AS, Trombitás K, Geach AJ, Yamasaki R, Sorimachi H, Granzier H, Gregorio CC, Labeit S. Myopalladin, a novel 145-kilodalton sarcomeric protein with multiple roles in Z-disc and I-band protein assemblies. Journal of Cell Biology. 2001;153:413–428.

12. Filomena MC, Yamamoto DL, Caremani M, Kadarla VK, Mastrototaro G, Serio S, Vydyanath A, Mutarelli M, Garofalo A, Pertici I. Myopalladin promotes muscle growth through modulation of the serum response factor pathway. Journal of Cachexia, Sarcopenia and Muscle. 2020;11:169–194.

13. Noureddine M, Gehmlich K. Structural and signaling proteins in the Z-disk and their role in cardiomyopathies. Frontiers in Physiology. 2023;14:1143858.

14. Meyer T, Ruppert V, Ackermann S, Richter A, Perrot A, Sperling SR, Posch MG, Maisch B, Pankuweit S. Novel mutations in the sarcomeric protein myopalladin in patients with dilated cardiomyopathy. European Journal of Human Genetics. 2013;21:294–300.

15. Beck MR, Dixon RD, Goicoechea SM, Murphy GS, Brungardt JG, Beam MT, Srinath P, Patel J, Mohiuddin J, Otey CA. Structure and function of palladin’s actin binding domain. Journal of molecular biology. 2013;425:3325–3337.

16. Sargent R, Liu DH, Yadav R, Glennenmeier D, Bradford C, Urbina N, Beck MR. Integrated structural model of the palladin–actin complex using XL-MS, docking, NMR, and SAXS. Protein Science. 2025;34:e70122.

17. Yadav R, Vattepu R, Beck MR. Phosphoinositide binding inhibits actin crosslinking and polymerization by palladin. Journal of molecular biology. 2016;428:4031–4047.

18. Abramson J, Adler J, Dunger J, Evans R, Green T, Pritzel A, Ronneberger O, Willmore L, Ballard AJ, Bambrick J. Accurate structure prediction of biomolecular interactions with AlphaFold 3. Nature. 2024;630:493–500.

19. Gurung R, Yadav R, Brungardt JG, Orlova A, Egelman EH, Beck MR. Actin polymerization is stimulated by actin cross-linking protein palladin. Biochem J. 2016;473:383–396. doi: 10.1042/BJ20151050

20. Heier JA, Dickinson DJ, Kwiatkowski AV. Measuring Protein Binding to F-actin by Co-sedimentation. JoVE (Journal of Visualized Experiments). 2017:e55613.

21. Amann KJ, Renley BA, Ervasti JM. A cluster of basic repeats in the dystrophin rod domain binds F-actin through an electrostatic interaction. Journal of Biological Chemistry. 1998;273:28419–28423.

22. Hüttelmaier S, Harbeck B, Steffens NO, Meßerschmidt T, Illenberger S, Jockusch BM. Characterization of the actin binding properties of the vasodilator-stimulated phosphoprotein VASP. FEBS letters. 1999;451:68–74.

23. Lee H-S, Bellin RM, Walker DL, Patel B, Powers P, Liu H, Garcia-Alvarez B, de Pereda JM, Liddington RC, Volkmann N. Characterization of an actin-binding site within the talin FERM domain. Journal of molecular biology. 2004;343:771–784.

24. Li X, Matsuoka Y, Bennett V. Adducin preferentially recruits spectrin to the fast growing ends of actin filaments in a complex requiring the MARCKS-related domain and a newly defined oligomerization domain. Journal of Biological Chemistry. 1998;273:19329–19338.

25. Tang JX, Szymanski PT, Janmey PA, Tao T. Electrostatic effects of smooth muscle calponin on actin assembly. European journal of biochemistry. 1997;247:432–440.

26. Dixon RD, Arneman DK, Rachlin AS, Sundaresan NR, Costello MJ, Campbell SL, Otey CA. Palladin is an actin cross-linking protein that uses immunoglobulin-like domains to bind filamentous actin. Journal of Biological Chemistry. 2008;283:6222–6231.

27. Bootman MD. Calcium signaling. Cold Spring Harbor perspectives in biology. 2012;4:a011171.

28. Dutta S, Tsiros C, Sundar SL, Athar H, Moore J, Nelson B, Gage MJ, Nishikawa K. Calcium increases titin N2A binding to F-actin and regulated thin filaments. Scientific reports. 2018;8:14575.

29. Greenfield NJ. Using circular dichroism spectra to estimate protein secondary structure. Nat Protoc. 2006;1:2876–2890. doi: 10.1038/nprot.2006.202

30. Bork P, Holm L, Sander C. The immunoglobulin fold: structural classification, sequence patterns and common core. Journal of molecular biology. 1994;242:309–320.

31. Otey CA, Dixon R, Stack C, Goicoechea SM. Cytoplasmic Ig-domain proteins: Cytoskeletal regulators with a role in human disease. Cell motility and the cytoskeleton. 2009;66:618–634.

32. Whitmore L, Wallace BA. Protein secondary structure analyses from circular dichroism spectroscopy: methods and reference databases. Biopolymers: Original Research on Biomolecules. 2008;89:392–400.

33. Miles AJ, Ramalli SG, Wallace BA. DichroWeb, a website for calculating protein secondary structure from circular dichroism spectroscopic data. Protein Sci. 2022;31:37–46. doi: 10.1002/pro.4153

34. Whitmore L, Wallace B. DICHROWEB, an online server for protein secondary structure analyses from circular dichroism spectroscopic data. Nucleic acids research. 2004;32:W668–W673.

35. Martinez-Martin I, Crousilles A, Ochoa JP, Velazquez-Carreras D, Mortensen SA, Herrero-Galan E, Delgado J, Dominguez F, Garcia-Pavia P, de Sancho D. Titin domains with reduced core hydrophobicity cause dilated cardiomyopathy. Cell Reports. 2023;42.

36. Mykkanen OM, Gronholm M, Ronty M, Lalowski M, Salmikangas P, Suila H, Carpen O. Characterization of human palladin, a microfilament-associated protein. Mol Biol Cell. 2001;12:3060–3073. doi: 10.1091/mbc.12.10.3060

37. Nakamura F, Osborn TM, Hartemink CA, Hartwig JH, Stossel TP. Structural basis of filamin A functions. J Cell Biol. 2007;179:1011–1025. doi: 10.1083/jcb.200707073

38. Tskhovrebova L, Trinick J. Titin: properties and family relationships. Nat Rev Mol Cell Biol. 2003;4:679–689. doi: 10.1038/nrm1198

39. Brand AH, Perrimon N. Targeted gene expression as a means of altering cell fates and generating dominant phenotypes. Development. 1993;118:401–415. doi: 10.1242/dev.118.2.401

40. Ranganayakulu G, Elliott DA, Harvey RP, Olson EN. Divergent roles for NK-2 class homeobox genes in cardiogenesis in flies and mice. Development. 1998;125:3037–3048. doi: 10.1242/dev.125.16.3037

41. Bang ML, Mudry RE, McElhinny AS, Trombitas K, Geach AJ, Yamasaki R, Sorimachi H, Granzier H, Gregorio CC, Labeit S. Myopalladin, a novel 145-kilodalton sarcomeric protein with multiple roles in Z-disc and I-band protein assemblies. J Cell Biol. 2001;153:413–427. doi: 10.1083/jcb.153.2.413

42. Gu Q, Mendsaikhan U, Khuchua Z, Jones BC, Lu L, Towbin JA, Xu B, Purevjav E. Dissection of Z-disc myopalladin gene network involved in the development of restrictive cardiomyopathy using system genetics approach. World journal of cardiology. 2017;9:320.

43. Auerbach D, Bantle S, Keller S, Hinderling V, Leu M, Ehler E, Perriard J-C. Different domains of the M-band protein myomesin are involved in myosin binding and M-band targeting. Molecular biology of the cell. 1999;10:1297–1308.

44. Linke WA. Stretching molecular springs: elasticity of titin filaments in vertebrate striated muscle. 2000.

45. Okagaki T, Weber FE, Fischman DA, Vaughan KT, Mikawa T, Reinach FC. The major myosin-binding domain of skeletal muscle MyBP-C (C protein) resides in the COOH-terminal, immunoglobulin C2 motif. The Journal of cell biology. 1993;123:619–626.

46. González-Morales N, Holenka TK, Schöck F. Filamin actin-binding and titin-binding fulfill distinct functions in Z-disc cohesion. PLoS Genetics. 2017;13:e1006880.

47. Benian GM, Mayans O. Titin and obscurin: giants holding hands and discovery of a new Ig domain subset. Journal of molecular biology. 2014;427:707.

48. Kostan J, Pavšič M, Puž V, Schwarz TC, Drepper F, Molt S, Graewert MA, Schreiner C, Sajko S, van der Ven PF. Molecular basis of F-actin regulation and sarcomere assembly via myotilin. PLoS biology. 2021;19:e3001148.

49. Gimona M, Djinovic-Carugo K, Kranewitter WJ, Winder SJ. Functional plasticity of CH domains. FEBS letters. 2002;513:98–106.

50. Paunola E, Mattila PK, Lappalainen P. WH2 domain: a small, versatile adapter for actin monomers. FEBS letters. 2002;513:92–97.

51. Way M, Pope B, Weeds A. Molecular biology of actin binding proteins: evidence for a common structural domain in the F-actin binding sites of gelsolin and α-actinin. Journal of Cell Science. 1991;1991:91–94.

52. Poukkula M, Kremneva E, Serlachius M, Lappalainen P. Actin-depolymerizing factor homology domain: a conserved fold performing diverse roles in cytoskeletal dynamics. Cytoskeleton. 2011;68:471–490.

53. Karp G. Cell and molecular biology: concepts and experiments. John Wiley & Sons; 2009.

54. Salmikangas P, van der Ven PF, Lalowski M, Taivainen A, Zhao F, Suila H, Schröder R, Lappalainen P, Fürst DO, Carpén O. Myotilin, the limb-girdle muscular dystrophy 1A (LGMD1A) protein, cross-links actin filaments and controls sarcomere assembly. Human molecular genetics. 2003;12:189–203.

55. Haywood NJ, Wolny M, Rogers B, Trinh CH, Shuping Y, Edwards TA, Peckham M. Hypertrophic cardiomyopathy mutations in the calponin-homology domain of ACTN2 affect actin binding and cardiomyocyte Z-disc incorporation. Biochemical Journal. 2016;473:2485–2493.

56. Angelini TE, Golestanian R, Coridan RH, Butler JC, Beraud A, Krisch M, Sinn H, Schweizer KS, Wong GC. Counterions between charged polymers exhibit liquid-like organization and dynamics. Proceedings of the National Academy of Sciences. 2006;103:7962–7967.

57. Tang JX, Janmey PA. The Polyelectrolyte Nature of F-actin and the Mechanism of Actin Bundle Formation (∗). Journal of Biological Chemistry. 1996;271:8556–8563.

58. Tang JX, Ito T, Tao T, Traub P, Janmey PA. Opposite effects of electrostatics and steric exclusion on bundle formation by F-actin and other filamentous polyelectrolytes. Biochemistry. 1997;36:12600–12607.

59. Cai L, Makhov AM, Bear JE. F-actin binding is essential for coronin 1B function in vivo. Journal of cell science. 2007;120:1779–1790.

60. Drmota Prebil S, Slapšak U, Pavšič M, Ilc G, Puž V, de Almeida Ribeiro E, Anrather D, Hartl M, Backman L, Plavec J. Structure and calcium-binding studies of calmodulin-like domain of human non-muscle α-actinin-1. Scientific reports. 2016;6:27383.

61. Nakamura F, Hartwig JH, Stossel TP, Szymanski PT. Ca2+ and calmodulin regulate the binding of filamin A to actin filaments. Journal of Biological Chemistry. 2005;280:32426–32433.

62. Yamamoto DL, Vitiello C, Zhang J, Gokhin DS, Castaldi A, Coulis G, Piaser F, Filomena MC, Eggenhuizen PJ, Kunderfranco P, et al. The nebulin SH3 domain is dispensable for normal skeletal muscle structure but is required for effective active load bearing in mouse. J Cell Sci. 2013;126:5477–5489. doi: 10.1242/jcs.137026

63. Spudich JA, Watt S. The regulation of rabbit skeletal muscle contraction: I. Biochemical studies of the interaction of the tropomyosin-troponin complex with actin and the proteolytic fragments of myosin. Journal of biological chemistry. 1971;246:4866–4871.

64. Albraiki S, Ajiboye O, Sargent R, Beck MR. Functional comparison of full-length palladin to isolated actin binding domain. Protein Science. 2023;32:e4638.

65. Zimmermann D, Morganthaler AN, Kovar DR, Suarez C. In vitro biochemical characterization of cytokinesis actin-binding proteins. Yeast Cytokinesis: Methods and Protocols. 2016:151–179.

66. Gasteiger E, Gattiker A, Hoogland C, Ivanyi I, Appel RD, Bairoch A. ExPASy: the proteomics server for in-depth protein knowledge and analysis. Nucleic acids research. 2003;31:3784–3788.

67. Sreerama N, Woody RW. Estimation of protein secondary structure from circular dichroism spectra: comparison of CONTIN, SELCON, and CDSSTR methods with an expanded reference set. Analytical biochemistry. 2000;287:252–260.

68. Andrade M, Chacon P, Merelo J, Morán F. Evaluation of secondary structure of proteins from UV circular dichroism spectra using an unsupervised learning neural network. *Protein Engineering*, Design and Selection. 1993;6:383–390.

69. Demeler B. Methods for the design and analysis of sedimentation velocity and sedimentation equilibrium experiments with proteins. Curr Protoc Protein Sci. 2010;Chapter 7:7 13 11-17 13 24. doi: 10.1002/0471140864.ps0713s60

